# Limb-localized induction of regeneration reshapes brain and amputation-site transcriptional states in adult *Xenopus laevis*

**DOI:** 10.64898/2026.10.23.761414

**Authors:** Stewart J. Russell, Kelsie A. Miller, David L. Kaplan, Michael Levin, Nirosha J. Murugan

**Author notes:** **Corresponding author:** Nirosha J. Murugan.

## Abstract

Limb regeneration requires coordinated tissue reconstruction and nervous system reintegration, yet whether regenerative induction alters brain responses to limb loss remains unclear. We analyzed transcriptomics from the paired brains and amputation sites of adult *Xenopus laevis* after hindlimb amputation with or without a 24-hour exposure to localized, pro-regenerative multi-drug treatment (MDT) delivered through a wearable bioreactor device. At 24 hours post treatment (24 HPT), MDT brains showed increased vascular and basement membrane transcripts and enrichment of oxidative phosphorylation and mTORC1 programs. In the limb, an exploratory integrated stress response (ISR) score increased at 11 and 24 HPT and returned to control levels by 7 days post treatment (7 DPT), coinciding with induction of growth differentiation factor 15 (GDF15), a cytokine responsive to mitochondrial stress. No corresponding increase in brain ISR score was detected. At 7 DPT, MDT brains showed enrichment of immune, heme metabolism and biosynthetic programs, reduced synaptic transmission gene expression and increased *gdf15.L*. Relative to an external uninjured reference, MDT and amputation-only brains showed concordant changes, with treatment modifying selected programs. Cross-species comparative analyses identified partial overlap with an axolotl regenerative glutamatergic signature and separation from selected mammalian injury profiles. These findings reveal cross-tissue responses to regenerative induction, identifying candidate mechanisms of brain–body communication in biomedically engineered vertebrate limb regeneration.

## Introduction

Regeneration of an amputated limb is more than a local problem of tissue replacement. A functional appendage must integrate patterned musculoskeletal tissue with vascular and peripheral nervous systems and re-enter the sensorimotor circuits that support sensation and movement. Amputation disrupts this integration by removing peripheral tissue, altering afferent input and changing central sensorimotor representations^1^. The reciprocal influence of injured limbs and the central nervous system has been described as the brain–limb axis^2^, yet it remains unknown whether the brain response to amputation changes when a local biomedical intervention redirects repair toward regeneration.

The nervous system is an essential component of the regenerative environment. In salamanders, denervation suppresses blastemal proliferation^3^, and arrests limb regeneration^4^, whereas a deviated nerve can support blastema formation and ectopic limb development when combined with appropriate positional information^5–7^. Hyperinnervation studies have also shown improved regenerative morphology in adult *Xenopus*, showing that nerve supply can modify the outcome in a species with restricted regenerative capacity^8^. Peripheral nerves therefore contribute trophic and patterning signals to the wound while providing a guiding scaffold between the limb and the central nervous system. While local functions in regeneration are well established, the consequences of regenerative activity for the brain are less well defined.

Limb loss has lasting consequences for brain organization and sensory experience. Phantom limb pain affects approximately 64% of amputees^9^, and loss of afferent input, through deafferentation-associated plasticity has been linked to changes in sensorimotor representations, gray matter and connectivity^10^. Early studies linked cortical reorganization to phantom pain^1,11^, although subsequent work showed that representations of missing hands can persist for years^12,13^. The central nervous systems changes may therefore involve altered regulation of established representations. Related plasticity occurs after peripheral nerve transection^14^ and surgical separation of fused digits^15^, with changes extending across sensorimotor networks^10,10^. Clinical interventions further connect peripheral innervation with positive sensory outcomes. Prosthesis use and restored sensory feedback have been associated with reduced phantom pain^16,17^. Agonist–antagonist myoneural interfaces preserve proprioceptive signaling and are associated with cortical proprioceptive activation comparable to that of non-amputees^18^, while targeted muscle reinnervation provides severed axons with a regenerative target and reduces pain^19^. Ablation of nociceptive neurons impairs wound healing and muscle regeneration in mice, and CGRP delivery restores both^20^, suggesting sensory afferents also contribute to repair. These observations link peripheral sensory input to cortical plasticity and tissue repair.

Evidence from regeneration-competent vertebrates suggest that appendage injury can engage responses beyond the amputation plane. In newts, limb amputation alters protein synthesis in the brain and in neural tissues on both sides of the spinal cord, while *Xenopus* hindlimb amputation induces rapid bioelectric responses in the contralateral uninjured limb^21–23^. In axolotls, adrenergic signaling couples local injury with systemic stem cell activation^24^. More recently, amputation was shown to activate a population of *etv1*-expressing glutamatergic neurons in the axolotl telencephalon that project to the hypothalamus, and inhibition of these neurons impaired tail regeneration^25^. Together, these studies show that appendage injury elicits distal neural and systemic responses.

Neural tissues are not the only means by which the state of an injured tissue can affect the brain. Cells also report their state systemically through circulating factors, many of which encode metabolic rather than sensory information^26,27^. Peripheral tissues can also communicate their metabolic state through circulating factors^28^. Regeneration imposes substantial energetic demands^29,30^, and the integrated stress response is a conserved cellular program that links disrupted homeostasis to transcriptional and secretory responses^31^. Growth differentiation factor 15 (GDF15) is induced by mitochondrial among other integrated stress pathways and can act as a secreted stress signal^31^. In mammals, circulating GDF15 acts through GFRAL receptors that are restricted to brainstem regions, regulating systemic energy balance^31–33^. GDF15 therefore provides a plausible molecular link between peripheral regenerative demand and the brain, although conservation of this signaling axis has not been established in adult *X. laevis*.

Adult *Xenopus laevis* provides a tractable system for distinguishing the response to limb loss from changes that accompany regenerative induction. Following hindlimb amputation, untreated adult frogs typically form a hypomorphic cartilaginous spike with limited pattern and function^34^. By contrast, an acute local intervention delivered through a wearable silk-hydrogel bioreactor, termed the BioDome, can redirect this outcome, in mouse and *Xenopus* models^35,36^. An initial progesterone treatment improved, but limited tissue patterning, innervation, vascular organization, and minimal locomotor activity^37^. Whereas a subsequent multidrug treatment, developed to stimulate the myriad of tissues within the hindlimb environment, induced durable, patterned, and functionally integrated limb regeneration with sensorimotor recovery over 18 months^38^. This matched injury model separates the transcriptional response to amputation alone from that associated with regenerative induction.

It remains unresolved whether such a local regenerative intervention also modifies the brain response to limb loss. Distinguishing a general response to trauma from adaptations that may support the construction and neural integration of a new appendage is important for identifying factors that reconnect the central nervous system with the regenerating periphery. It may also reveal intended or unintended system-level consequences of interventions designed to act locally. We therefore used paired whole-brain and amputation-site RNA sequencing from the same animals to determine whether a 24-hour limb-localized pro-regenerative treatment produces a brain response distinct from amputation alone, across across early and later timepoints. We also compared the frog brain response with published regenerative competent, axolotl and selected non-regenerative mouse pain and injury datasets to distinguish features associated with regenerative induction from those shared with peripheral injury more broadly. MDT was associated with temporally structured transcriptional remodeling across brain and limb, including an early limb ISR-associated program, changes in brain metabolic and neural gene expression and delayed induction of brain *gdf15*. These findings identify the brain as part of the systemic response accompanying induced limb regeneration and establish candidate pathways for further functional testing.

## Results

### Limb-localized regenerative treatment alters brain and amputation-site transcriptional trajectories

Appendage amputation initiates a local wound response while also altering physiology at sites distant from the injury^21,23,24^. A 24-hour multidrug treatment (MDT) delivered to the stump in a wearable BioDome redirects adult *Xenopus laevis* toward prolonged, patterned outgrowth^38^. We therefore asked whether this limb-localized intervention also altered the brain transcriptional response to amputation. We analyzed paired bulk RNA sequencing profiles from whole brain and amputation-site tissue collected from the same animals after unilateral hindlimb amputation. Animals received MDT in the BioDome or underwent amputation without a device (ND; Fig. 1a–c). Brain was collected at 24 HPT and 7 DPT, and matched limb tissue was collected at the same intervals, with an additional limb sample at 11 HPT (Fig. 1b–d). Amputation-site tissue is termed blastema where needed for consistency with the source dataset. The MDT versus ND comparison therefore captures the combined device and drug condition relative to amputation alone.

**Figure 1.**
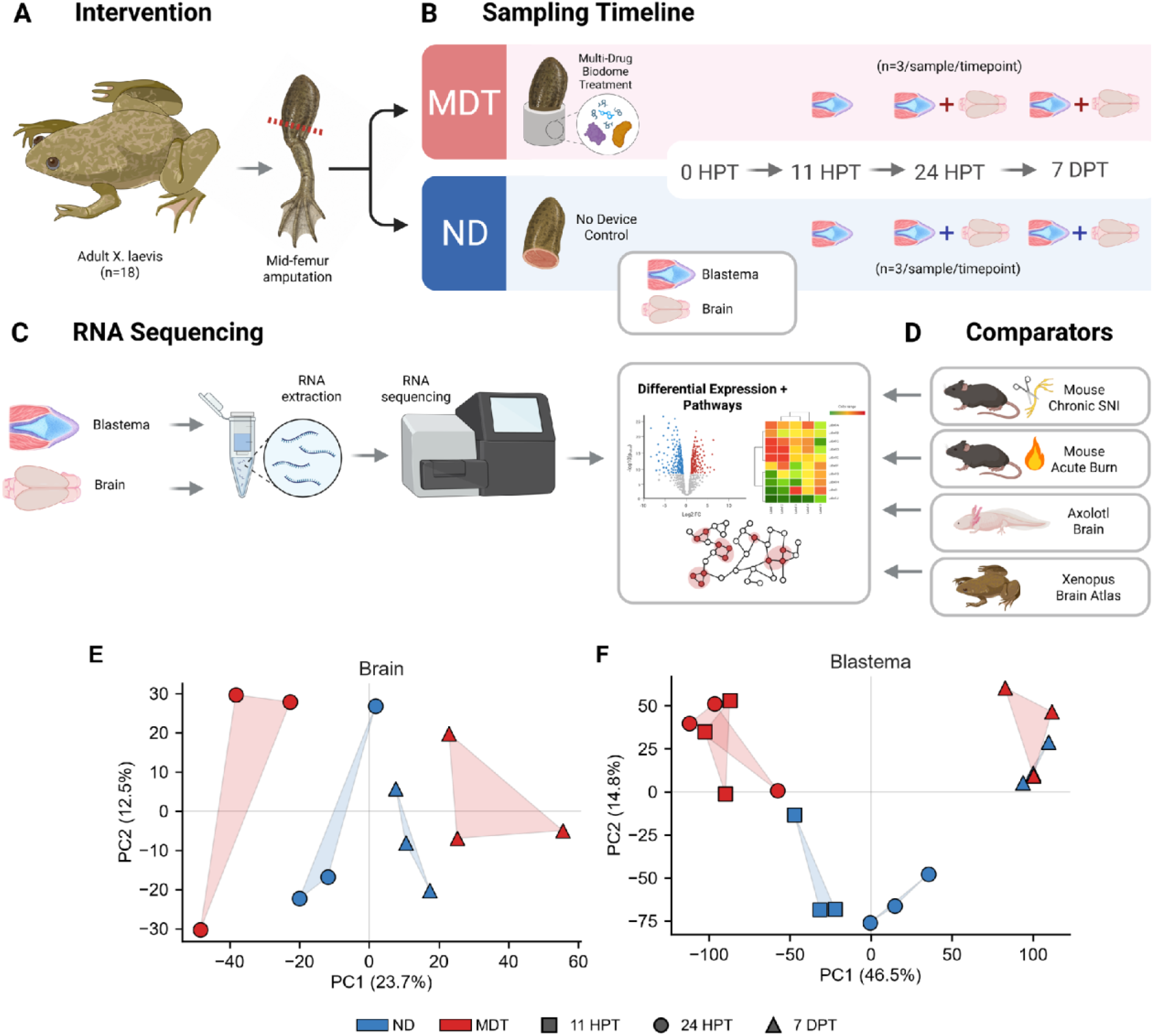
Experimental design, tissue sampling and global transcriptional structure. **(A)** Adult *Xenopus laevis* received a hindlimb amputation at the midpoint of the tibiofibular bone and were assigned to multidrug Biodome treatment (MDT) or a no-device control (ND). **(B)** Brain and blastema were sampled in pairs across a shared time course, given in hours and days post-treatment (HPT, DPT): brain at 24 HPT and 7 DPT, and blastema at 11 HPT, 24 HPT and 7 DPT (n = 3 per time point and condition). **(C)** Both tissues underwent bulk RNA sequencing followed by differential expression and pathway analysis. **(D)** External comparator datasets included mouse chronic spared-nerve injury, mouse cortical responses to acute peripheral injury (burn and formalin), an axolotl brain single-cell atlas and a *Xenopus* brain atlas. **(E, F)** Principal component analysis (PCA) of **(E)** brain and **(F)** amputation-site samples. Color denotes condition (ND blue, MDT red) and symbol denotes time point, with axis labels giving the variance explained by each component. Panels A–D created in BioRender.

Principal component analysis revealed treatment- and time-associated structure in the brain transcriptomes, with PC1 and PC2 accounting for 23.7% and 12.5% of the total variance, respectively (Fig. 1e). Condition-associated separation was more pronounced in the brain than in the matched amputation-site samples (Fig. 1f). These global patterns motivated direct testing of treatment and temporal comparisons.

Direct MDT and ND comparisons identified 29 differentially expressed genes in the brain at 24 HPT and 32 at 7 DPT (adjusted *P* < 0.05; Fig. 2a). Within MDT brains, 932 genes differed between 24 HPT and 7 DPT, compared with 9 genes across the same interval in ND brains (Fig. 2b–d). This threshold-based comparison indicated a substantially larger temporal shift in the treated group, although the difference in DEG counts does not itself constitute a formal treatment-by-time interaction test. Matched blastemas also showed a treatment-associated regenerative profile marked by lower expression of fibrosis-related ECM and collagen-synthesis programs (Supplementary Figs. 1 and 2). Together, the MDT BioDome condition was associated with local limb-tissue remodeling and a temporally structured brain response.

**Figure 2.**
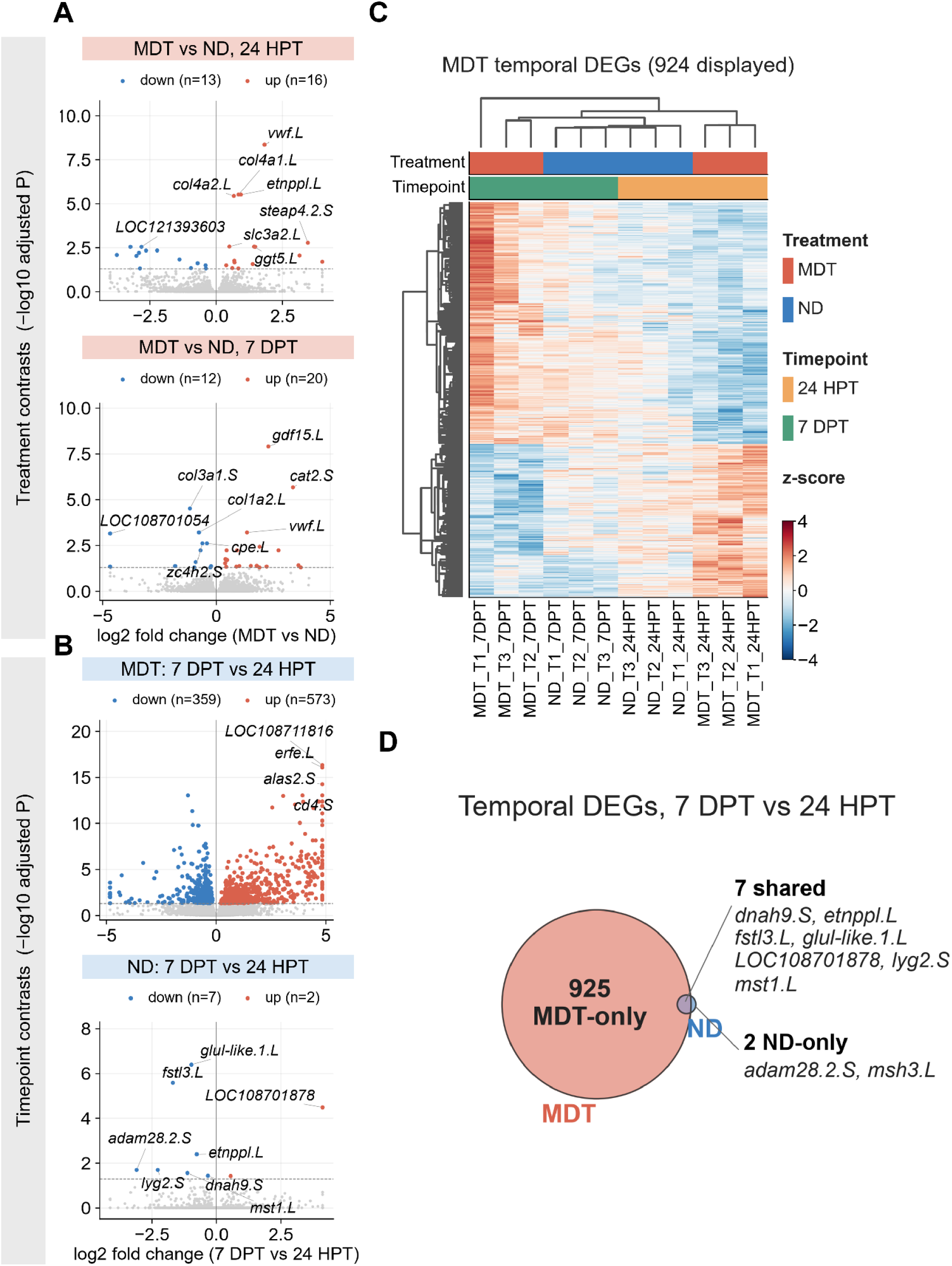
MDT brain transcriptomes show greater temporal change. **(A)** Volcano plots of brain differential expression, MDT versus ND, at 24 HPT (top) and 7 DPT (bottom); colored points are differentially expressed genes (up in MDT, red; down in MDT, blue) and selected genes are labeled. The suffixes .L and .S denote the L and S homeologs of the *X. laevis* allotetraploid genome. **(B)** Within-condition volcano plots of the 7 DPT versus 24 HPT contrast, for MDT (top) and ND (bottom). **(C)** Unsupervised clustering of the MDT-temporal differentially expressed genes (DEGs) across all brain samples; color is row z-scored expression, and the Treatment and Timepoint bars annotate the columns. **(D)** Overlap of the 7 DPT versus 24 HPT DEG sets between the two treatment conditions.

### Early treatment-associated brain responses are enriched for neurovascular, basement membrane, and metabolic programs

Vascular and extracellular matrix (ECM) remodeling help establish a permissive wound environment and provide structural and positional information during appendage regeneration^39,40^. We used gene-set enrichment analysis (GSEA) to define the programs that distinguished MDT and ND brains at 24 HPA. Relative to ND brains, MDT brains showed positive enrichment of the Hallmark epithelial-to-mesenchymal transition gene set (NES = +1.56), oxidative phosphorylation (NES = +1.45) and mTORC1 signaling (NES = +1.41; adjusted *P* < 0.05; Fig. 3a,c). In this context, the epithelial-to-mesenchymal transition annotation was driven primarily by matrix and vascular genes and is best interpreted as an ECM-associated transcriptional program.

**Figure 3.**
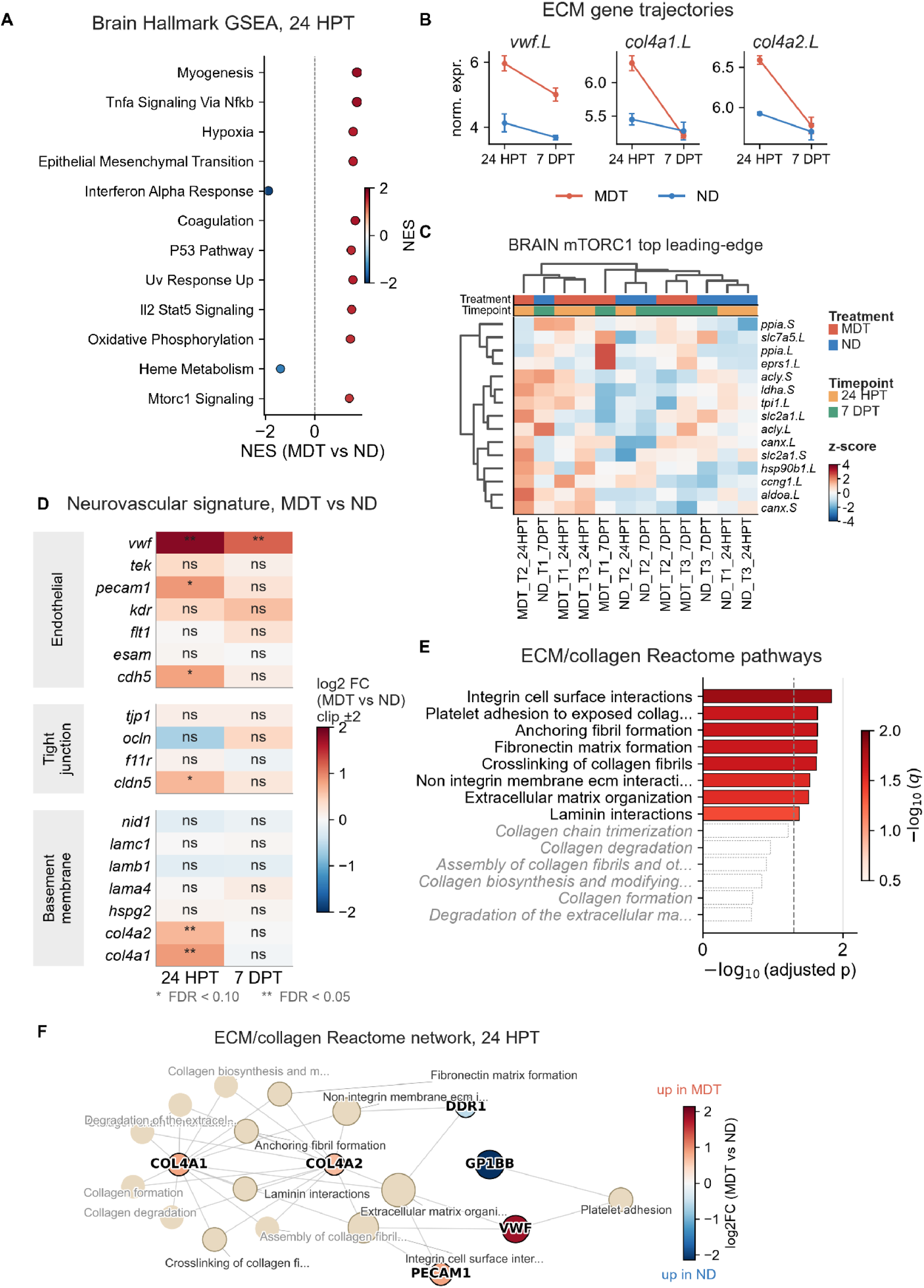
Early (24 HPT) brain response shows vascular, ECM and metabolic gene enrichment. **(A)** Hallmark GSEA, MDT vs ND at 24 HPT; x-axis/color is NES, dot size reflects significance. **(B)** Normalized expression trajectories of *vwf.L, col4a1.L, col4a2.L*; line connects group means, error bars s.e.m. **(C)** Heatmap of leading-edge gene expression for mTORC1 pathway genes (color, row z-score). **(D)** Neurovascular signature (endothelial, tight-junction, basement-membrane genes); color is MDT-vs-ND log2 fold change per timepoint; asterisks denote significance, ns = not significant. **(E)** ECM/collagen Reactome over-representation; filled bars significant, open not. **(F)** Gene-pathway network; gene nodes colored by log2 fold change, significant pathway nodes outlined.

Differentially expressed genes included vascular, endothelial, and basement membrane-associated factors. *vwf* increased in MDT brains (log2 fold change=+1.8, adjusted P=4.4×10^-9^), while *col4a1* and *col4a2* were also differentially expressed (adjusted P<1×10^-5^; Fig. 3b,d). Changes in *cldn5*, together with *vwf*, *col4a1*, and *col4a2*, were consistent with a vascular-associated transcriptional response.

To further define the processes represented by the early differentially expressed genes, we performed Reactome over-representation analysis which identified 22 significant terms among genes expressed more highly in MDT brains, whereas one term was significant among genes expressed less highly (adjusted P<0.05). Enriched terms included integrin cell-surface interactions (4 of 20 genes, adjusted P=0.0016), anchoring fibril formation, fibronectin matrix formation, and collagen fibril crosslinking (adjusted P=0.013 for each; Fig. 3e, f). These terms were largely driven by a common set of genes, including *col4a1*, *col4a2*, *vwf*, *pecam1*, *cdh5*, and *gja1*, indicating a coherent response rather than multiple independent pathways. The collagen genes were type IV basement-membrane collagens typical of healthy brain tissue composition rather than the fibrillar collagens associated with scar formation. Overall, the early MDT brain phenotype was characterized by coordinated vascular, basement-membrane, oxidative phosphorylation, and mTORC1-associated gene expression.

### The brain response transitions toward immune-metabolic and biosynthetic regulation by 7 days

Appendage regeneration proceeds through temporally ordered phases of wound repair, blastema formation, and tissue outgrowth which often occur over the course of days^39,40^. We therefore examined whether the early brain programs persisted at 7 DPT during the early regenerative response. The treatment-associated brain response at this later timepoint was dominated by immune and metabolic Hallmark programs. Heme metabolism (NES=+2.09), allograft rejection (NES=+2.00), interferon-gamma response (IFN-γ) (NES=+1.87), interferon-alpha (IFN-α) response (NES=+1.78), and MYC targets (NES=+1.79) were positively enriched in MDT relative to ND brains (adjusted P<0.002 for all; Fig. 4a). Heme metabolism showed the strongest enrichment and concordant changes across its leading-edge genes (adjusted P=3.0×10^-11^; Fig. 4b).

**Figure 4.**
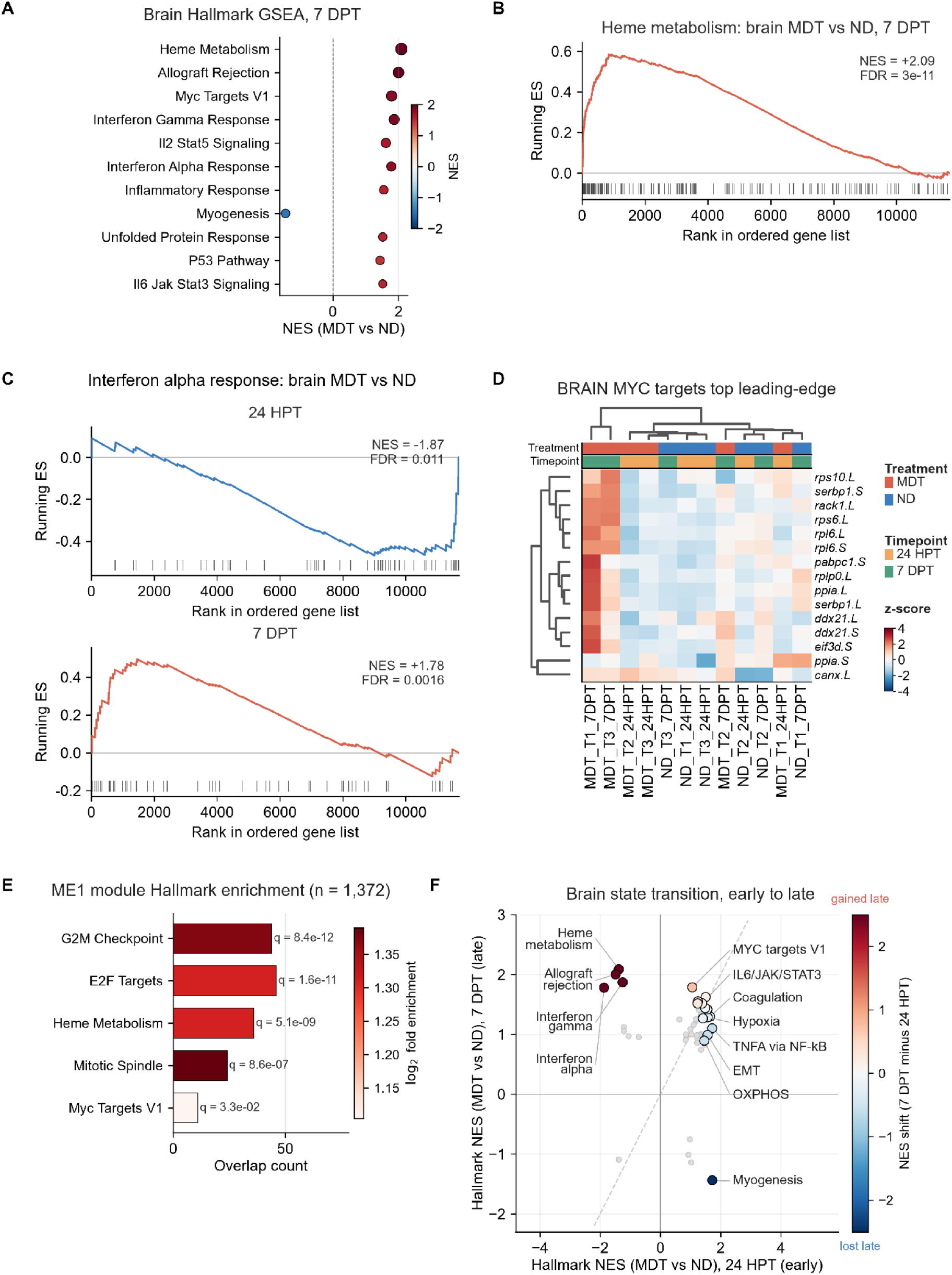
Late (7 DPT) brain response dispplayed immune, heme and biosynthetic gene-set enrichment. **(A)** Hallmark GSEA, MDT versus ND at 7 DPT; NES on the x-axis and color, dot size reflects significance. **(B)** Running-enrichment (GSEA) plot for the heme metabolism Hallmark, MDT versus ND at 7 DPT. **(C)** Running-enrichment plots for the interferon-α response at 24 HPT and 7 DPT. **(D)** Leading-edge expression heatmap for MYC targets (color, row z-score). **(E)** Hallmark overrepresentation within the treatment-associated co-expression module (turquoise); bar length is the number of overlapping genes and color is fold enrichment. **(F)** Per-Hallmark NES at 24 HPT versus 7 DPT, colored by the shift from early to late, separating an early vascular-metabolic from a late immune-metabolic state.

The IFN-α response was negatively enriched at 24 HPT (NES=−1.87, adjusted P=0.011; Fig. 3a) and positively enriched at 7 DPT (NES=+1.78, adjusted P=0.0016; Fig. 4c). MYC-target and protein-biosynthesis programs were also elevated at 7 DPT, supported by concordant changes in leading-edge ribosomal and translation-associated genes (NES=+1.79, adjusted P=3.3×10^-6^; Fig. 4d). These patterns indicate a temporal reorganization of immune and biosynthetic gene expression rather than a uniform increase or decrease in inflammatory signaling.

To determine whether the same biological programs would emerge from the data without reference to curated gene sets, we constructed brain co-expression networks using Weighted Gene Co-expression Network Analysis (WGCNA). The largest module (module 1, named turquoise) showed a treatment-by-time interaction in its eigengene (β=+0.44, P=0.043). This module was enriched for G2M checkpoint (44 genes, adjusted p=8.4×10^-12^), E2F targets (46 genes, adjusted p=1.6×10^-11^), heme metabolism (36 genes, adjusted p=5.1×10^-9^), and MYC targets V1 (11 genes, adjusted P=0.033; Fig. 4e). Thus, an unsupervised analysis independently recovered the programs identified by gene-set enrichment. Comparison of the MDT-versus-ND enrichment profiles at the two timepoints showed a shift from early vascular, ECM, and metabolic programs toward later immune-metabolic, cell-cycle, and biosynthetic programs (Fig. 4f).

### GDF15 emerges as a candidate transcriptional signal linking the limb and brain

To identify candidate mediators of the paired brain and blastema response, we examined the genes that most strongly distinguished MDT and ND animals. At 7 DPT, *gdf15* was the most significant of the 32 differentially expressed genes within the brain. In mammals, GDF15 is secreted by stressed peripheral tissues and acts through GFRAL, a receptor with restricted expression in the hindbrain^33^. This established peripheral-to-central signaling function made GDF15 a candidate for cross-tissue communication during regenerative induction.

Brain *gdf15.L* expression did not reach the differential-expression threshold between MDT and ND animals at 24 HPT (log2 fold change=+0.87, adjusted P=0.79). By 7 DPT, however, *gdf15.L* was higher in MDT brains (log2 fold change=+2.28, adjusted P=1.3×10^-8^), with a significant treatment-by-time interaction (β=+1.41, P=0.0044; Fig. 5a). In the limb tissue, *gdf15.L* was already elevated at 24 HPT (log2 fold change=+3.7, adjusted p=0.04; Fig. 5b). Across brain samples, *gdf15.L* expression correlated with the eigengene of the largest treatment-associated co-expression module (module 1, turquoise; Spearman rho=+0.70, P=0.011; Fig. 5c), linking its expression to the broader late brain program.

**Figure 5.**
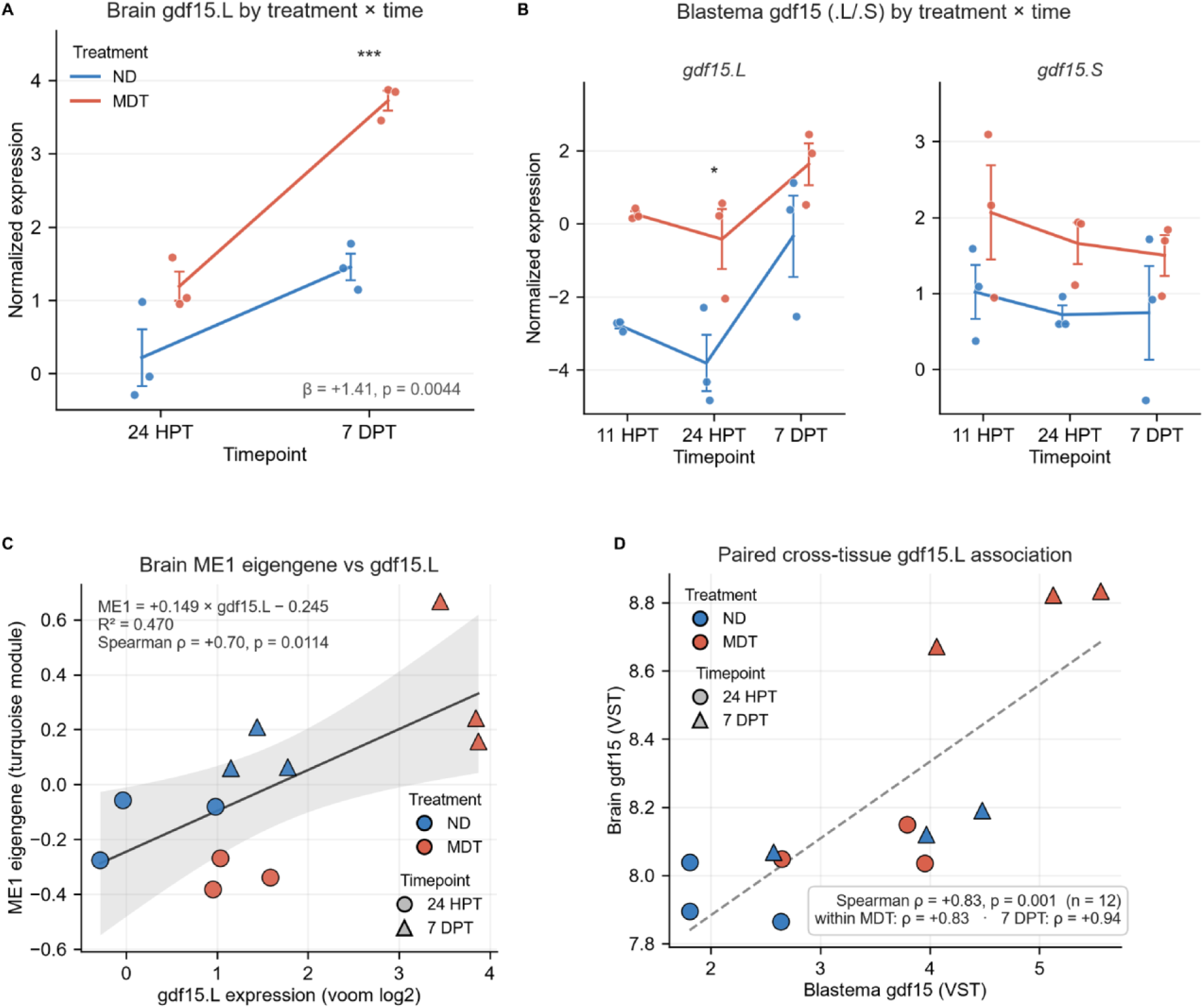
*gdf*15 expression covaries between brain and amputation-site tissue. **(A)** Brain *gdf15.L* expression by condition and timepoint. Lines connect group means and error bars represent SEM. **(B)** Amputation-site *gdf15.L* (left) and *gdf15*.S (right) expression by condition and timepoint. **(C)** Brain turquoise module eigengene (ME) against *gdf15.L* expression, with a linear fit and its 95% confidence band; color indicates condition and symbol indicates timepoint. **(D)** Paired per-animal association between amputation-site and brain *gdf15.L* expression (variance-stabilized); color indicates condition, symbol indicates timepoint and the dashed line is a linear fit. In A and B, asterisks denote significant MDT-versus-ND differences.

Since brain and blastema were collected from the same animals, we next investigated whether *gdf15.L* expression covaried across tissues. Brain and blastema expression showed a strong positive correlation across the 12 paired samples (Spearman rho=+0.83, P=0.001; Fig. 5d). The association was strongest at 7 DPT (rho=+0.94) and within MDT animals (rho=+0.83) and was weaker within ND animals (rho=+0.64) and at 24 HPT (rho=+0.38). The pooled correlation combines treatment and time-point groups and therefore does not establish direct coupling or the direction of signaling (peripheral-to-central vs central-to-peripheral). Nevertheless, the delayed increase in brain *gdf15.L*, its association with the major treatment-responsive brain module, and its covariation with paired blastema expression identify *gdf15* as a candidate cross-tissue mediator for functional testing.

The integrated stress response (ISR) is a conserved intracellular program that integrates disruptions to cellular homeostasis and can induce secreted stress signals including GDF15^31^. We tested whether the early limb *gdf15* increase occurred within a broader ISR-associated response. Using an exploratory *Xenopus* adaptation of a human ISR index, we excluded GDF15 from the primary score and retained 101 of 119 genes in the limb. Scores were higher in MDT animals at 11 HPA (MDT minus ND = +1.40, adjusted *P* = 0.0076) and 24 HPA (+1.96, adjusted *P* = 0.0014), with little treatment separation at 7 DPA (+0.07, adjusted *P* = 0.923; Supplementary Fig. 3). The treatment-by-time analysis supported attenuation between 24 HPA and 7 DPA (difference in treatment effects = −1.88, adjusted *P* = 0.010). Both *atf3* homeologs and *ddit3.L* were also upregulated in limb tissue at 24 HPA (genome-wide adjusted *P* ≤ 1.7 × 10^−8^). No corresponding increase in the adapted brain ISR score was detected at either interval. Early limb *gdf15* induction therefore coincided with a transient ISR-associated program. The later brain increase occurred without detectable elevation of the adapted brain ISR score.

### Synaptic transmission gene expression declines later in the regenerative trajectory

Loss of peripheral input and nerve injury can induce persistent structural and transcriptional changes in central neural circuits^1,41^ with pain as a common clinical correlate. In regeneration-competent axolotls, appendage injury also engages an *etv1*/*slc17a7*-positive glutamatergic brain population required for efficient tail regeneration^25^. We therefore asked whether MDT altered neural transcriptional programs relative to amputation alone. At 7 DPT, MDT brains showed negative enrichment of synaptic-transmission gene sets, including regulation of trans-synaptic signaling (NES = −2.10, adjusted *P* = 1.5 × 10^−6^; Fig. 6a). Eighteen synaptic- and axon-associated Gene Ontology terms were negatively enriched at 7 DPT, whereas none reached significance at 24 HPT. At the sampled intervals, this treatment-associated difference was detected at 7 DPT and was not detected at 24 HPT.

**Figure 6.**
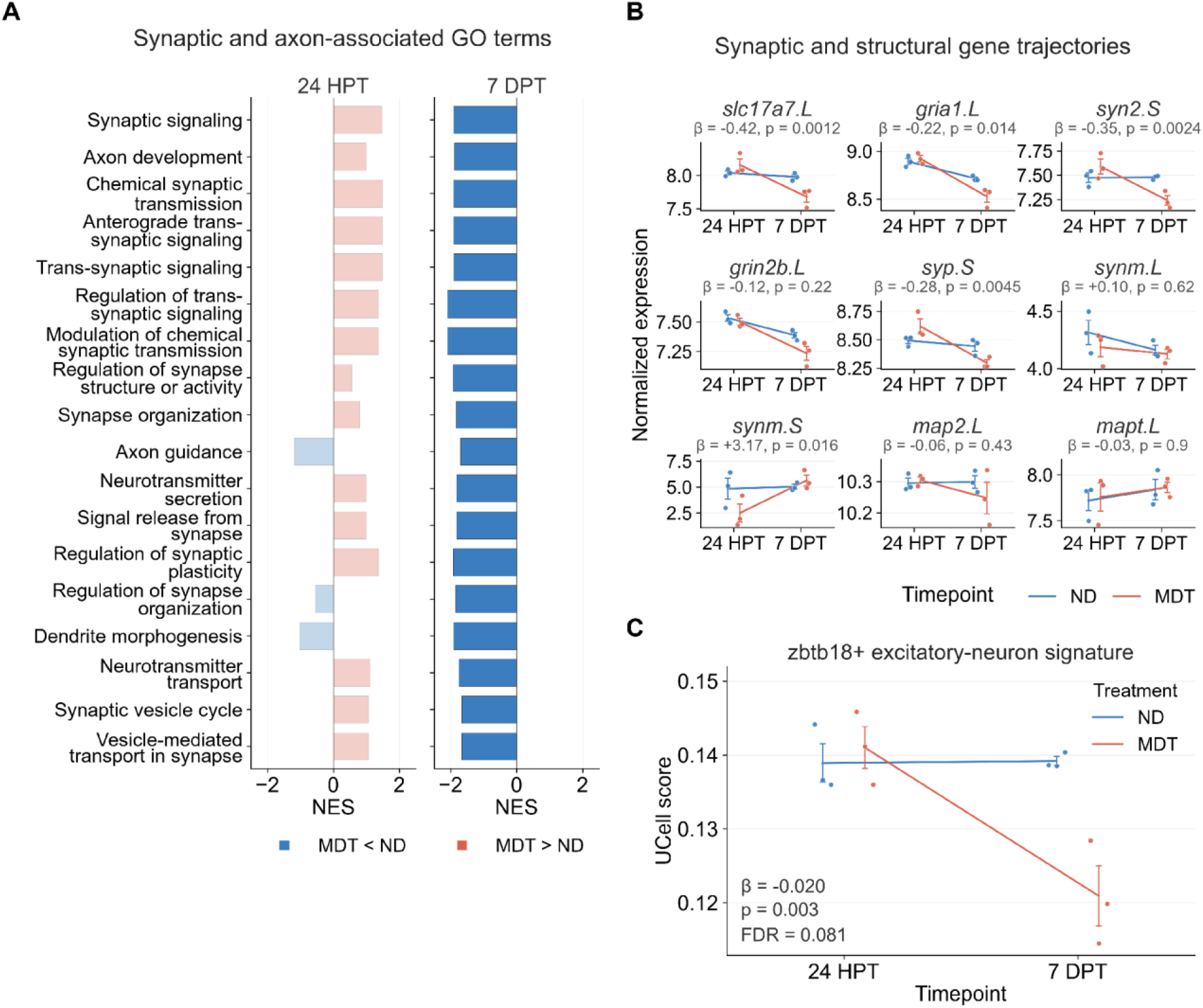
Decreased expression of synaptic transmission-associated transcripts in MDT-treated brains at 7 DPT. **(A)** Synaptic and axon-associated Gene Ontology biological-process (GO:BP) term enrichment at 24 HPT and 7 DPT, MDT versus ND; bar length shows the NES, color indicates the direction of change (MDT lower than ND, blue; MDT higher than ND, coral) and bars are faded where not significant. **(B)** Expression trajectories of synaptic and structural genes across timepoints; points are individual animals, lines connect group means and error bars show s.e.m. **(C)** Excitatory-neuron (*zbtb18*+) signature score (UCell) in brain across timepoints, by condition.

To determine whether the bulk transcriptional signal aligned with a defined neural cell-type program, we projected signatures from an independent adult *X. laevis* single-cell brain atlas onto the bulk transcriptomes^42^. The largest nominal decrease was observed for a *zbtb18*-positive excitatory-neuron signature (β = −0.020, nominal *P* = 0.003; Fig. 6c), although this association did not remain significant after correction across the 26 cell-type signatures tested (adjusted *P* = 0.081). Genes contributing to the synaptic signal included *slc17a7*, *gria1*, *syn2* and *syp*, with a treatment-by-time interaction for *slc17a7* (*P* = 0.0012; Fig. 6b). These data support lower expression of synaptic-transmission-associated programs in MDT brains at 7 DPT. Bulk RNA sequencing cannot distinguish per-cell transcriptional changes from altered cellular composition and cannot infer neuronal activity.

### Pro-regenerative treatment, MDT, alters the magnitude and timing of a shared post-amputation brain state

Both MDT and ND animals underwent limb amputation, and the primary experiment did not include uninjured brain controls. Because appendage loss can evoke molecular and bioelectric responses at sites distant from the wound^21,23,24^, treatment-associated differences must be interpreted within a broader post-amputation state. To resolve the temporal organization of the treatment effect, we first summarized higher-order biological processes at each time point. This analysis captured late enrichment of protein biosynthesis, endocrine, immune, and heme-associated programs, together with late suppression of synaptic and cholesterol-associated expression (Fig. 7a).

**Figure 7.**
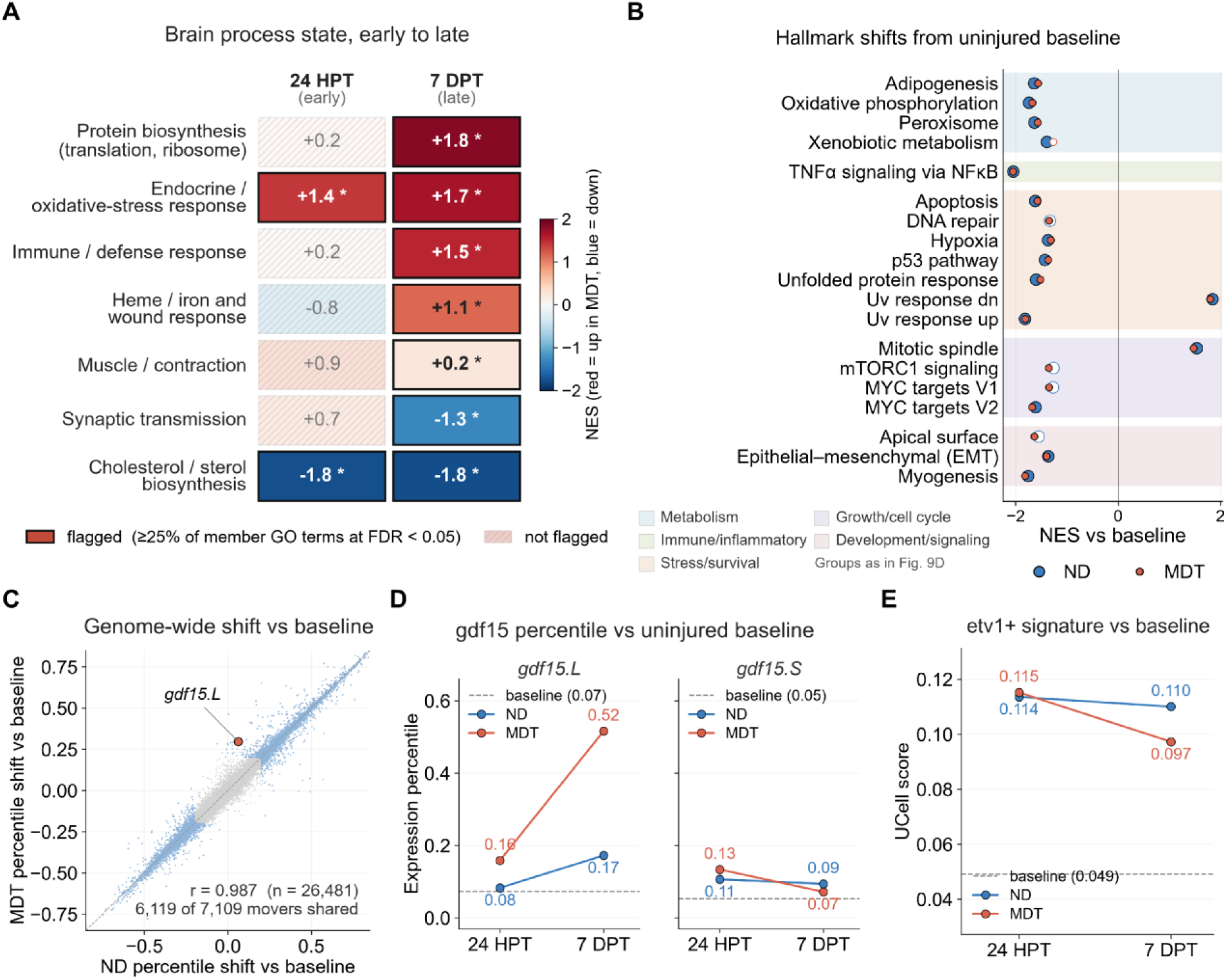
Brain process-state timeline and comparison with an uninjured baseline. **(A)** Seven GO:BP process clusters scored at 24 HPT (early) and 7 DPT (late); tile color shows the mean member NES (coral higher, blue lower in MDT) and a star flags clusters in which at least 25% of member terms reach FDR<0.05. **(B, C, D, E)** Comparison of each condition against an uninjured *X. laevis* brain baseline. **(B)** Hallmark percentile-rank shift from baseline in each condition; filled symbols indicate FDR<0.05 within that condition and open symbols do not. Rows are grouped by broad biological function for display; the groups are descriptive and are not separately tested. **(C)** Genome-wide concordance of the two conditions’ shifts from baseline, with *gdf15.L* highlighted as a leading off-diagonal gene. **(D)** *gdf15.L* (left) and *gdf15*.S (right) expression percentile within each dataset, by condition and timepoint, relative to the uninjured baseline (dashed line). (**E**) Score (UCell) of the axolotl *etv1*+ regeneration-associated signature (Fig. 9A, B) in brain, by condition and timepoint, relative to the uninjured baseline (dashed line).

We then used an independent uninjured adult *X. laevis* brain dataset as a qualitative baseline (n=2 brains, GSE73419;^43^). Because these samples were generated in a separate experiment, no differential expression statistics were calculated across datasets. Relative to this baseline, MDT and ND brains shifted in the same direction across nearly all Hallmark programs (Fig. 7b), and their genome-wide rank shifts were highly concordant (r=0.987; Fig. 7c). These findings indicate that a substantial component of the brain response was shared by both amputated groups.

Against this common post-amputation state, *gdf15.L* remained one of the strongest genes separating MDT and ND brains. Its expression increased further from the uninjured baseline in MDT animals, whereas the *gdf15.S* homeolog remained stable (Fig. 7c, d). Both treatment conditions also exhibited the axolotl *etv1*+ regeneration-associated signature at approximately 2.3-fold the uninjured level at 24 HPT. By 7 DPT, this signature decreased to 2.0-fold in MDT brains and remained at 2.2-fold in ND brains (^25^; Fig. 7e). Together, these comparisons suggested that MDT modified the magnitude and timing of selected transcriptional programs within a response largely shared after amputation.

### Paired brain and limb transcriptomes show treatment-associated covariation

The paired expression of *gdf15.L* suggested that treatment-associated variation may likewise be coordinated across tissues. To determine whether this relationship extended beyond a single candidate gene, we integrated the matched brain and blastema transcriptomes using multi-omics factor analysis (MOFA). The analysis identified one cross-tissue factor associated with the treatment-by-time interaction (Factor 3, adjusted P=0.029). Factor 3 accounted for 11.6% of the variance in the blastema and 3.3% of the variance in the brain (Fig. 8a).

**Figure 8.**
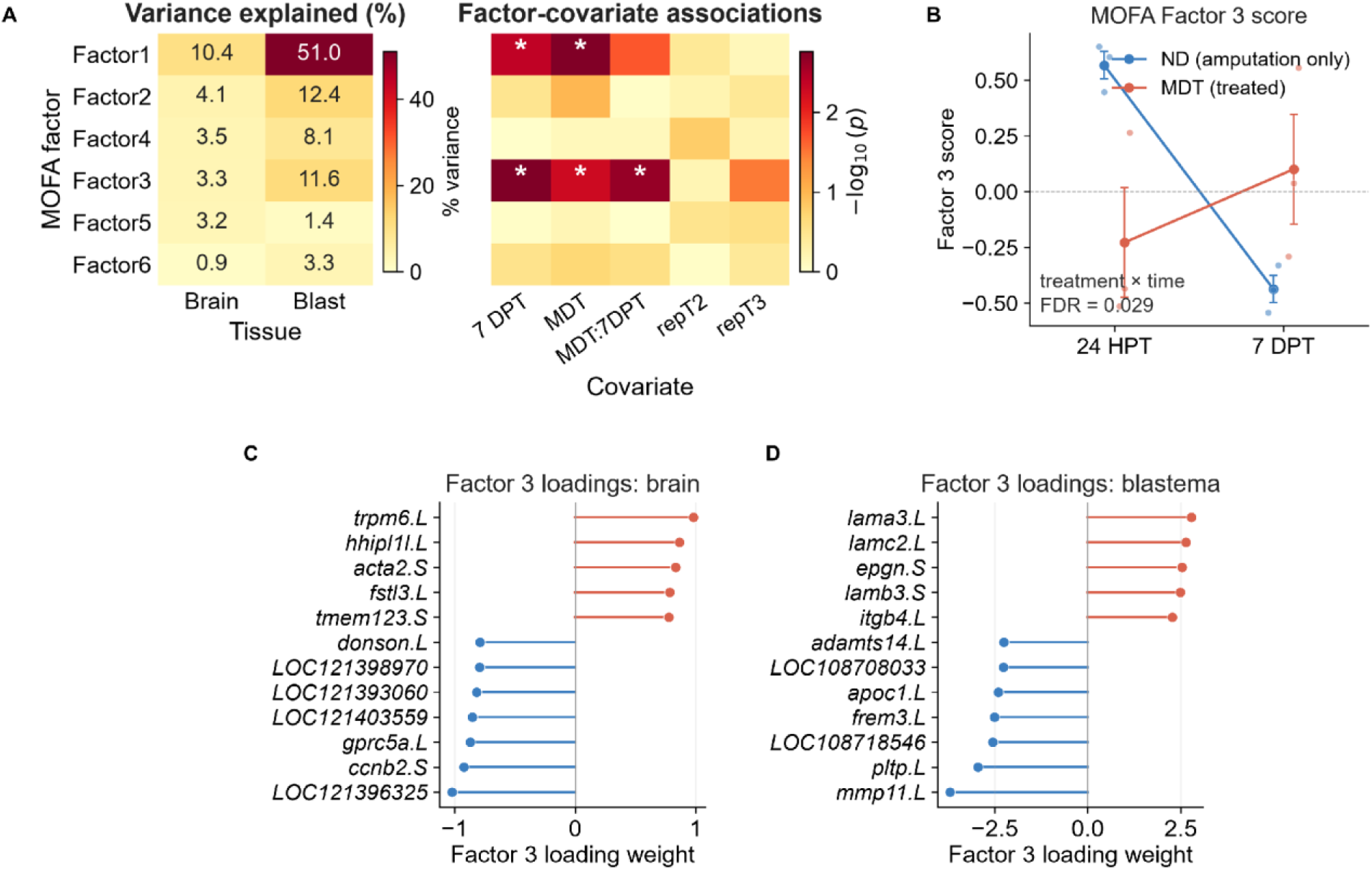
Multi-omics factor analysis links the paired brain and blastema. **(A)** Multi-omics factor analysis (MOFA) variance explained per factor and tissue (left) and factor-covariate associations (right); asterisks mark significant associations; repT2 and repT3 are replicate terms (replicate 2 or 3 versus 1). **(B)** Factor 3 scores by condition and timepoint. **(C, D)** Top Factor 3 loadings for the **(C)** brain and **(D)** blastema views.

MDT and ND animals followed opposing temporal directions along Factor 3, between 24 HPT and 7 DPT (Fig. 8b). In the brain, the factor loadings captured cell-cycle, transport, and structural-remodeling-associated genes (Fig. 8c). In the blastema, the same factor captured ECM and regenerative-remodeling-associated genes (Fig. 8d). This latent factor demonstrates coordinated transcriptional covariation across the paired tissues. It does not establish whether the limb influences the brain, the brain influences the limb, or both tissues respond to a shared systemic signal.

### Cross-species comparisons place the brain response in regenerative and injury contexts

To determine whether the MDT-associated brain response resembled neural programs described after regenerative or non-regenerative injury, we compared it with published axolotl and mouse datasets. In axolotl, an *etv1*-positive glutamatergic brain population responds to appendage injury and is required for efficient tail regeneration^25^. After mapping this signature to frog orthologs, it was detected in both *X. laevis* treatment conditions and showed a more negative temporal change in MDT than in ND brains (treatment-by-time β = −0.028, *P* = 0.024; Fig. 9c). These data identify partial transcriptional overlap with an axolotl regeneration-associated signature. Projection of a small cross-species marker set onto whole-brain RNA sequencing cannot establish a homologous frog cell population.

**Figure 9.**
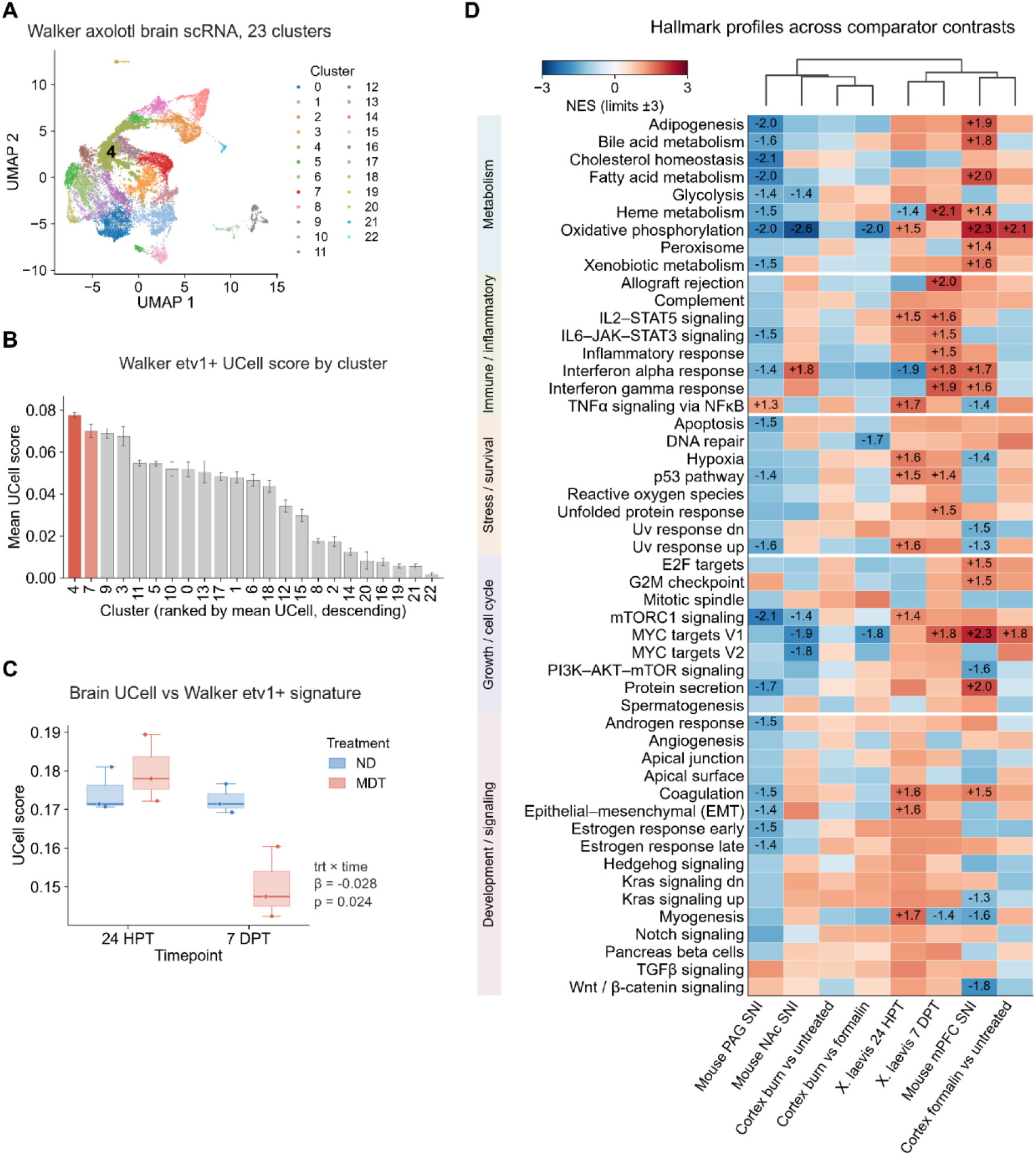
Cross-species comparison of the MDT-associated brain response. **(A)** UMAP of the Walker axolotl brain single-cell atlas. Clusters are color-coded and the *etv1*-positive glutamatergic cluster is enlarged. **(B)** *etv1*-positive glutamatergic signature score per cluster. **(C)** Axolotl *etv1*-positive signature score in *X. laevis* brain. **(D)** Hallmark NES across mouse chronic spared-nerve injury in periaqueductal gray (PAG), nucleus accumbens (NAc) and medial prefrontal cortex (mPFC), mouse acute cortical injury and the two *X. laevis* contrasts. Columns were ordered by Ward’s hierarchical clustering of the full Hallmark NES profiles. Axolotl data are shown in A–C and were not included in D.

For mammalian comparisons, we analyzed mouse spared-nerve-injury datasets from the periaqueductal gray, nucleus accumbens and medial prefrontal cortex^25^, together with acute hind-paw burn and formalin datasets from primary somatosensory cortex^44^. Across 50 Hallmark gene sets, the two frog MDT-versus-ND profiles clustered were adjacent in hierarchical clustering and separated from the selected chronic-pain and acute-injury profiles (Fig. 9d). Several immune and metabolic gene sets were shared across datasets. The complete MDT-associated profile did not match any single mouse comparison. These analyses are descriptive because the datasets differ in species, brain region, sampling interval, injury paradigm and comparison design, and the acute mouse comparisons contain only one library per condition. The observed separation cannot be attributed uniquely to regeneration.

## Discussion

Regenerative competence is conventionally defined by events at the injured tissue, including resolution of the wound response, suppression of fibrosis, establishment of progenitor states, and reactivation of developmental programs capable of reconstructing missing anatomy. Our findings expand that view by showing that induction of a pro-regenerative trajectory in the adult *X. laevis* hindlimb is accompanied by a distinct and time-dependent transcriptional state in the brain. Paired analysis further identified treatment-associated covariation between brain and blastema (Fig. 8), indicating that the physiological state established during regenerative induction extends beyond the amputation plane. Together with evidence for systemic injury responses in regenerative vertebrates, including contralateral bioelectric signaling, systemic cell cycle activation, long-range adrenergic coordination and brain-dependent regenerative control, these data support a systemic physiology of appendage regeneration^23–25,45^.

A particularly informative feature of this response was its temporal patterning, whereby metabolic stress responses were temporally ordered across tissues. The brains of treated animals were enriched in the early phase for oxidative phosphorylation and mTORC1 signaling together with vascular and basement-membrane-associated transcripts (Fig. 3), whereas the later state shifted toward heme metabolism, MYC targets, protein biosynthesis, cell cycle and immune programs (Fig. 4). The limb ISR score increased at 11 and 24 HPT and returned to control levels by 7 DPT, while the early brain response was enriched for oxidative phosphorylation and mTORC1-associated expression (Supplementary Fig. 3; Fig. 3). This temporal patterning places metabolic adaptation early in the response to regenerative induction and is consistent with experimental work showing that bioenergetic state can help determine regenerative competence^46^. Oxygen sensing, HIF1A stability, glycolytic regulation and mitochondrial state have emerged as determinants of whether injured vertebrate tissues access regenerative cell states, linking metabolism to cell plasticity and regenerative outcome^29,30^. In this context, the metabolic programs observed in the brain enrichment raise the possibility that the energetic demands of regenerative reconstruction at the wound site become represented systemically. Regeneration requires coordinated proliferation, protein synthesis, extracellular-matrix turnover, vascular growth and redox adaptation^30,39^. Bioenergetic state may therefore function as biological information that reports the physiological demands of reconstruction across tissues.

The early brain response also contained a coherent neurovascular expression. Early treatment-associated transcripts included *vwf*, *pecam1*, *cdh5*, *cldn5*, *col4a1* and *col4a2*, together with enrichment of integrin interactions, fibronectin matrix organization and basement-membrane-associated processes (Fig. 3b-f). COL4A1 and COL4A2 are type IV basement-membrane collagens and, together with endothelial junctional genes, are consistent with a neurovascular-associated transcriptional response rather than a fibrillar-collagen scar signature. This distinction is particularly relevant because the treated blastema simultaneously shows reduced fibrosis-associated extracellular matrix and collagen programs as it enters a pro-regenerative trajectory (Supplementary Figs. 1 and 2). Neural and vascular systems are tightly coupled during development and repair, with endothelial cells, neurons, glia and extracellular matrix sharing guidance and trophic mechanisms while vascular state regulates oxygen delivery, metabolic support and access to circulating signals^47^. Vascular and basement-membrane transcripts were therefore among the earliest treatment-associated changes in the brain.

The ISR analysis provided a separate measure of cellular stress. The ISR is a conserved cell-intrinsic program that translates disrupted homeostasis into transcriptional and secretory responses^31^. Its transient elevation in MDT limb tissue at 11 and 24 HPT places the early limb *gdf15* increase within a broader stress-associated program that had subsided by 7 DPT (Supplementary Fig. 3). No corresponding increase in the adapted brain ISR score was detected, including at 7 DPT when brain *gdf15.L* was elevated. The later brain increase therefore cannot be interpreted as evidence of a brain-wide ISR, and GDF15 expression alone is not specific evidence of mitochondrial energetic stress. In mammals, stress-induced GDF15 can signal through GFRAL-expressing hindbrain neurons^31–33,48^, and macrophage-derived GDF15 supports inflammatory resolution and regeneration after muscle injury^49^. The early limb induction, later brain expression, association with the treatment-responsive brain module and covariance across paired tissues make GDF15 a candidate for functional testing in *Xenopus* (Fig. 5). The present data do not show that limb-derived GDF15 protein entered the circulation, reached the brain or activated a conserved receptor pathway. Brain *gdf15.L* may have been induced locally, and the pooled cross-tissue correlation may partly reflect treatment and time.

The delayed (7 day) neuronal response reveals a second dimension of this distributed regenerative state. By 7 days, synaptic transmission-associated programs were broadly shifted downward in treated brains, with contributions from genes spanning glutamatergic transmission and synaptic organization, including *slc17a7*, *gria1*, *syn2* and *syp* (Fig. 6a,b). Projection of an independent *Xenopus* single-cell atlas associated this effect most strongly with a *zbtb18*+ excitatory-neuron signature (Fig. 6c;^42^). This pattern is consistent with altered excitatory-neuronal state during regenerative progression. Sustained excitatory drive can recruit slower homeostatic programs that reduce excitatory synaptic strength after earlier changes in intrinsic excitability^50^, providing a plausible explanation for why a synaptic signature emerges later rather than immediately after amputation. Mature presynaptic machinery can also constrain neuronal growth competence, with reduction of vesicle-priming components increasing axonal regenerative capacity after adult CNS injury^51^. Attenuation of mature synaptic programs may therefore mark a period of enhanced neural plasticity in which established circuit states are loosened while the nervous system adapts to altered afferent input and a regenerating appendage.

The axolotl comparison places this neural-state change into a regenerative context. A discrete population of *etv1*+/*slc17a7*+ glutamatergic neurons in the axolotl medial pallium was identified that responds to appendage injury and contributes functionally to regeneration through hypothalamic neurotensin and neuroendocrine signaling^25^. In our cross-species analysis, the corresponding regeneration-associated neuronal signature was detectable in the *Xenopus* brain following amputation in both treatment groups and followed a different later trajectory after pro-regenerative treatment (Figs. 7e and 9a-c). This suggests that adult *Xenopus* retains elements of a CNS injury-response program used by a highly regenerative vertebrate. Regenerative competence may therefore depend partly on how conserved neuronal programs are temporally sequenced, maintained and resolved after injury. The later reduction in synaptic-associated expression in treated animals may represent progression through an activated injury-responsive state toward a different phase of neural plasticity.

The selected mouse datasets provided a separate reference for brain responses to non-regenerative injury and chronic pain. Chronic spared-nerve injury in mice produces broad transcriptional remodeling in brain regions linked to pain, affect and stress, including the medial prefrontal cortex, periaqueductal gray and nucleus accumbens^52^. Acute burn and formalin injury also alter cortical transcription, although on a shorter timescale and with a smaller pathway-level response^44^. Across Hallmark gene sets, the *Xenopus* treatment contrasts clustered together and remained separated from these mammalian injury states (Fig. 9d). The distinction matters because BioDome-treated animals ultimately recover sensorimotor function^38^. These comparisons do not exclude pain-related contributions: both frog groups were amputated, MDT may alter nociception, and the studies differ in species, brain region, sampling time, and experimental contrast.

This has direct implications for the eventual functional reintegration of a regenerated appendage. Limb loss changes afferent input and can produce persistent alterations in sensorimotor representations and connectivity^1,11–13^. A biologically regenerated limb must ultimately reconnect with a nervous system that has already adapted to its absence. A limb replacement or regenerative therapy may therefore need to preserve, restore, or re-engage the neural representation of that limb so that the regenerated appendage can become functionally reintegrated into the body. Work with prostheses, brain–machine interfaces and robotic augmentation shows that the adult brain readily builds and remodels representations of artificial effectors that replace or extend a limb, and that cortical network connectivity reorganizes even after chronic amputation^53–55^. Regenerative induction and neural interfaces may thus be two routes by which a changed periphery drives adaptive central reorganization, and the treated-brain state described here may represent a regenerative analogue of the extensive central remodeling that follows amputation in mammals^10^. Reconstructive approaches that preserve physiological proprioceptive signaling support this principle by maintaining cortical responses closer to those of intact limbs, while sensory neurons themselves can regulate inflammatory and regenerative processes in peripheral tissues through neuroimmune signaling^18,20^. Neural reintegration should therefore be considered part of regenerative outcome. The present transcriptomes cannot determine whether the MDT-associated brain state facilitates later sensorimotor recovery, but they identify processes and time points at which that relationship can be tested.

Taken together, the data support a distributed transcriptional response to regenerative induction. Local positional information, extracellular matrix composition, innervation, and developmental signaling remain fundamental determinants of pattern formation, while metabolic, vascular, endocrine, bioelectric and neural states add further dimensions through which the organism can coordinate its response to tissue loss^6,23–25^. In this view, the regenerating limb, brain and other tissues participate in a shared physiological state whose trajectory may influence regenerative outcome. The cross-tissue latent factor identified here provides transcriptomic support for such a distributed state (Fig. 8).

Several limitations define the current resolution of this study. Whole-brain bulk transcriptomics cannot localize the vascular, immune, metabolic and neuronal programs to specific nuclei or cell populations, and the absence of perfusion leaves a possible contribution from retained blood. The external uninjured dataset provides a qualitative reference rather than an experimentally matched baseline. Because MDT is the intervention that induces the regenerative outcome, the design cannot separate an acute treatment response from molecular processes required for subsequent regeneration or from direct systemic effects of compounds released from the BioDome. The absence of a profiled device-only condition also prevents separation of drug effects from device effects. Cross-tissue covariance does not establish direct communication or its direction. Direct measurement of circulating GDF15, mapping of *gfral* and *ret* expression and perturbation of the ligand-receptor pathway would test whether a conserved endocrine mechanism operates in adult *Xenopus*. Spatial and functional interrogation of candidate neuronal populations, paired with direct measures of limb and brain metabolism, will be required to distinguish central responses that report regenerative state from those that contribute to it.

Collectively, these findings expand the spatial and physiological domain in which engineered regeneration should be considered. Regenerative induction in the adult *Xenopus* limb is accompanied by a transition in brain state that begins with neurovascular and bioenergetic programs, progresses toward immune, biosynthetic and neuronal reorganization, and covaries with molecular changes in the regenerating appendage (Figs. 3-8). GDF15 provides a candidate link between regenerative bioenergetic demand and long-range signaling, while the axolotl and mammalian comparator analyses place the treated brain state within a broader landscape of regenerative and injury-associated neural responses (Fig. 9). The relevant unit of analysis for regenerative engineering may therefore be larger than the wound itself. Testing whether the brain state we describe causally contributes to regenerative outcome represents a logical next step toward the integration of the brain-limb axis into therapy design.

## Methods

### Animals and multidrug treatment

The tissues and sequencing data were generated in the experiment described by Murugan et al.^38^; no new animal procedures were performed for this reanalysis. The original handling and experimental procedures were approved by the Tufts University Institutional Animal Care and Use Committee, as reported in the original study. Eighteen adult female *Xenopus laevis* received a right hindlimb amputation through the midpoint of the femur. Nine animals received a wearable silk-hydrogel bioreactor (BioDome) loaded with a five-component pro-regenerative multidrug treatment (MDT: BDNF, growth hormone, 1,4-DPCA, retinoic acid and resolvin D5) applied to the stump 1 hour after amputation and maintained for 24 hours; the other nine received amputation alone with no device (ND). The original device-only condition was not profiled here. Analysis files, code and Supplementary Data retain the original study’s CT label for MDT and its hours/days-post-amputation (HPA/DPA) labels for the same collection points.

### Tissue collection and RNA sequencing

Blastema at the amputation site was collected at 11, 24 and 168 hours post-treatment (HPT, counted from removal of the device; 168 HPT = 7 days post-treatment, DPT), and whole brain at 24 HPT and 7 DPT, with three biological replicates per treatment, timepoint and tissue. Brain and blastema were sampled from the same animals at the two shared timepoints, giving a paired cross-tissue design. Libraries were prepared with the TruSeq Stranded RNA kit with Ribo-Zero Gold rRNA depletion and sequenced on an Illumina HiSeq 2500 as single-end 51-nt reads, yielding 30 libraries (12 brain, 18 blastema).

### Read processing and quantification

Reads were adapter- and quality-trimmed with fastp v0.24.0 (Phred≥20, minimum length 36;^56^) and checked with FastQC before and after trimming^57^. Trimmed reads were aligned to the *X. laevis* v10.1 assembly (XENLA_10.1, Xenbase, NCBI RefSeq annotation;^58^) with STAR v2.7.11b in two-pass mode^59^, giving approximately 82% uniquely mapped reads across brain libraries. Because the libraries are reverse-stranded, gene-level counts were taken from the reverse-strand column of the STAR ReadsPerGene output and aggregated into a gene-by-sample matrix. Salmon v1.10.2 was additionally used to quantify transcript abundance against a decoy-aware Xenbase index^60,61^ for the cross-dataset baseline comparison described below.

### Filtering and normalization

Genes above 1 count per million in at least three samples were retained (edgeR;^62^) and TMM normalization factors were calculated within each tissue for the limma-voom analyses^63^. A variance-stabilizing transform of the filtered counts (VST; DESeq2, blind=TRUE;^64^) was used for the co-expression, factor and cell-type signature analyses described below. Per-tissue principal-component analysis (Fig. 1e,f) used limma-voom log2 counts-per-million of homeolog-summed counts (.L and .S summed per gene) for the 5,000 most variable genes. Expression trajectories and expression heatmaps display per-sample TMM-normalized log2 counts-per-million from limma-voom (^65,66^; row-scaled to z-scores in heatmaps), with significance annotations taken from the DESeq2 contrasts described below. The cross-tissue *gdf15* coupling analysis used VST values.

### Differential expression

Differential expression was computed per tissue with DESeq2 (design=∼0+group, where group encodes each treatment-timepoint combination), Wald tests with independent filtering (alpha=0.05) and a Benjamini-Hochberg (BH) adjusted P<0.05 threshold^67^, with no fold-change cutoff. The .L and .S homeologs of the allotetraploid genome were kept separate; per-homeolog statistics are primary for all gene-level results. Brain contrasts were MDT versus ND at each timepoint and the within-condition temporal contrasts (7 DPT versus 24 HPT in each condition); the temporal-DEG heatmap and Venn overlap intersect the two per-condition temporal DEG sets. Blastema contrasts comprised MDT versus ND and the within-condition temporal comparisons across all three timepoints. Full results for all 13 contrasts are provided in Supplementary Data 1. A homeolog-summed count matrix (counts of .L and .S summed per gene) was analyzed identically to provide pooled statistics for over-representation testing. Brain *gdf15* analyses used *gdf15.L*, the dominant brain-expressed homeolog. Because brain and blastema derive from the same animals, per-animal cross-tissue coupling was quantified as the Spearman correlation of paired brain and blastema *gdf15.L* values across the 12 matched samples, with within-MDT, 7-DPT-only and homeolog-summed correlations as robustness checks. For the curated neurovascular signature panel, MDT-versus-ND log2 fold changes are shown per timepoint with significance from a within-signature limma test on homeolog-collapsed genes (within-signature FDR<0.10 *, <0.05 **).

### Ortholog mapping, gene-set enrichment and over-representation

For Hallmark and Reactome analyses, *X. laevis* genes were mapped to human orthologs by removing the .L/.S suffix and matching upper-cased symbols. Where both homeologs mapped to the same ortholog, the first occurrence in the significance-ordered results was retained, so each ortholog contributes exactly one value. Genes were ranked by sign(log2 fold change) ×-log10(P). Hallmark enrichment shown in the dot plots was computed with the GSEA interface of clusterProfiler^68^, which wraps the fgsea algorithm, against MSigDB collections retrieved with msigdbr^69,70^. Running-enrichment plots and the process-timeline and baseline analyses used fgsea directly (minSize=15, maxSize=500, eps=0;^71^). The synaptic Gene Ontology Biological Process (GO:BP) panels used clusterProfiler gseGO on the same signed ranking with *X. laevis* gene-level GO annotation (org.Xl.eg.db;^72^); the GO analyses therefore test different term universes after size filtering (1,756 terms for the gseGO panels, 3,473 for the process timeline), and FDR values are internal to each analysis. Gene sets at BH-adjusted P<0.05 were considered enriched, and normalized enrichment scores (NES) are reported throughout; leading-edge heatmaps show row-scaled expression of the leading-edge genes of the named pathway. For the brain at 24 HPT, Reactome over-representation used clusterProfiler::enricher^73^ on genes at BH-adjusted P<0.1 in the homeolog-summed MDT-versus-ND contrast, and the matrix and collagen pathways were displayed as a gene-concept network with gene nodes colored by log2 fold change.

### Co-expression networks (WGCNA)

Signed weighted co-expression networks were built per tissue from VST expression of the top 8,000 most variable genes (WGCNA;^74^), with the soft-thresholding power chosen by scale-free-topology fit (a fixed power of 14 where the estimate was undefined) and modules detected with blockwiseModules (signed TOM, minModuleSize=30, mergeCutHeight=0.25). Module eigengene effects were tested per module with linear models (eigengene ∼ treatment × time), and the dominant treatment-associated brain module (module 1, turquoise) was characterized by Hallmark over-representation of its member genes and by Spearman correlation of its eigengene with *gdf15.L*.

### Multi-omic factor analysis (MOFA+)

Coordinated brain–blastema variation was modeled with MOFA+^75^. The input was the 12 animals with matched brain and blastema (three per condition and timepoint), represented as two views (brain and blastema). Each view contained VST expression of its top 8,000 most variable genes and was scaled to unit variance. The model used Gaussian likelihoods and up to 10 factors; factors explaining less than 1% of variance were dropped. Each retained factor’s scores were tested with a linear model (score ∼ treatment × time + replicate) and BH-adjusted across factor-term combinations. Factor 3 was the only factor associated with the treatment-by-time interaction and is reported with its variance explained per tissue and its top loadings (top 200 genes by absolute weight per view).

### Brain process-state timeline

GO:BP fgsea (as above) was computed for the MDT-versus-ND contrast at 24 HPT and 7 DPT across 3,473 terms. The 193 terms reaching FDR<0.05 at either timepoint were grouped by average-linkage hierarchical clustering of Wang semantic similarity (GOSemSim;^76,77^) into twelve clusters; seven biologically coherent clusters are displayed, with two closely related synaptic clusters merged. Each process is plotted as the mean NES of its member terms per timepoint and flagged when at least 25% of member terms individually reach FDR<0.05; no separate process-cluster significance test was performed.

### Cell-type signature projection (Xenopus brain atlas)

From the adult *X. laevis* single-cell brain atlas (GSE195790;^42^), each annotated cell type with≥30 cells was summarized as its top 50 marker genes by log fold change of CPM-normalized cell-type pseudobulk against the mean of other cell types, with no ortholog mapping required (same species). Signatures were scored on each bulk brain library (VST expression) with UCell^78^ and tested with linear models (score ∼ treatment × time + replicate), BH-adjusted across the 26 testable cell-type signatures.

### Axolotl single-cell comparator

Published axolotl (*Ambystoma mexicanum*) brain single-cell RNA-seq spanning uninjured, tail-amputation, limb-amputation and muscimol conditions (^25^; SRR31800874-SRR31800877) was quantified against the AmexT_v47 transcriptome with Salmon alevin^79^ and imported with tximport^80^. Cells with≥500 genes, ≥1,000 UMIs and below 25% mitochondrial reads were retained, normalized with SCTransform^81^, integrated across samples by canonical-correlation anchors (Seurat;^82^), reduced by PCA (30 components) and UMAP, and clustered at resolution 0.6; clusters were annotated with published marker genes scored by UCell. Enrichment of the *etv1*+ regeneration-associated cluster across conditions was tested with Fisher exact tests. The *etv1*+ signature was scored on the *X. laevis* bulk brain samples with UCell after upper-case symbol matching; the signature comprises 13 *X. laevis* homeologs of seven of the eight published markers (*ntng1* has no resolved homeolog).

### Mammalian injury comparators

Mouse spared-nerve-injury (SNI) bulk RNA-seq of medial prefrontal cortex, nucleus accumbens and periaqueductal gray (GSE91396;^52^) was analyzed from the deposited RPKM matrix (GEOquery;^83^): log2(RPKM+1) values were tested for SNI-versus-sham differential expression with limma using an intensity-trend prior, followed by Hallmark GSEA (clusterProfiler; *Mus musculus* msigdbr). Acute-injury contrasts in primary somatosensory cortex sampled 1 hour post-injury (hind-paw burn versus untreated, formalin versus untreated, and burn versus formalin; PRJNA1137110;^44^) were quantified with Salmon against the mouse GENCODE vM33 transcriptome and imported with tximport. With one library per condition, fold changes are descriptive log2 counts-per-million ratios without replicate-based statistics; genes were ranked by log2 fold change for the same Hallmark GSEA, and these contrasts are interpreted at pathway level only. The comparator heatmap shows NES for all 50 Hallmark gene sets across eight contrasts (the two *X. laevis* MDT-versus-ND contrasts and the six mouse contrasts), with columns ordered by Ward hierarchical clustering and NES values labeled where BH-adjusted P<0.05.

### Uninjured baseline anchor

Because the primary experiment includes no uninjured condition, an external uninjured adult *X. laevis* whole-brain dataset (GSE73419, two biological replicates from independent clutches;^43^) was used as a qualitative, rank-based anchor; no differential-expression statistics were computed between datasets. The Murugan and GSE73419 libraries were quantified together with Salmon (TPM) and restricted to protein-coding genes detected at TPM≥1 in any replicate of either dataset (27,478 genes), mitigating the library-preparation (poly-A versus Ribo-Zero) and read-layout (paired-end 2×100 versus single-end 51) differences. For each gene, an expression percentile was computed within each library. A gene’s rank shift is its percentile in an amputated condition minus its percentile in the uninjured baseline, giving one genome-wide shift vector per condition. Agreement between the two conditions was summarized as the Pearson correlation of their shift vectors. Each condition’s shift vector was also used as the ranking metric for pre-ranked fgsea against the Hallmark and Reactome collections, identifying the programs that shifted relative to baseline. Cross-dataset validity was checked with paired GSE73419 eye samples: photoreceptor and phototransduction markers were strongly eye-enriched while housekeeping genes were equivalent across tissues. All baseline panels are reported as qualitative anchors given the n=2 baseline and the strain, sex, age and batch differences between studies.

### Statistics and reproducibility

Downstream analyses used R 4.5.0 with Bioconductor 3.21 (R Core Team, 2025;^84^) and the packages cited above, with msigdbr 26.1.0^85^. Multiple testing was controlled by the BH procedure within each analysis at the thresholds stated above. Stochastic steps (fgsea, MOFA+ training, Seurat integration and clustering) used fixed random seeds (42, 20260430 and 20260501, respectively).

### Use of AI tools

Large language model-based assistants (Claude, Anthropic; Codex, OpenAI) were used to write and run analysis and figure-generation code under the direction and review of S.J.R., and to help revise and edit the text. The authors checked all code, results, figures and text and take full responsibility for the content.

### Exploratory integrated stress response scoring

The human ISR–GDF15 index of Smith et al.^31^ was adapted using curated Xenbase human-ortholog assignments. TMM-normalized log2 counts-per-million were averaged across homeologs assigned to one human symbol and z-standardized per gene within each tissue.

GDF15 was excluded from the primary score, leaving 101/119 genes in limb and 97/119 in brain. For each observed gene subset, regression scoring coefficients were reconstructed as the inverse human reference correlation matrix multiplied by the published GDF15-associated factor loadings, using the 339 fibroblast training samples and rounded loadings supplied with the published code. The adapted score was the weighted sum of *Xenopus* gene z-scores; its absolute scale was not compared between tissues. Ordinary linear models of animal-level scores used treatment–timepoint groups within each tissue (three independent animals per group), with two-sided t-tests for treatment, temporal and interaction contrasts. BH adjustment covered 119 tests in the exploratory ISR family: 17 contrasts across both tissues for six score variants and a canonical ISR target-gene average. Sensitivity analyses varied GDF15 inclusion, reference-SD scaling and loading-based weights; using the higher-expressed homeolog was analyzed as a separate 119-test family. All tests and the adapted scoring coefficients are provided in Supplementary Data 2. The ISR analysis used R 4.6.1 and limma 3.68.5, without empirical-Bayes moderation or gene-level precision weights for the composite scores.

## Data availability

Raw sequencing reads and processed count matrices will be deposited at GEO/SRA on publication. Differential expression results for all contrasts are provided in Supplementary Data 1. Comparator datasets are available under their original accessions: Walker et al. 2025 (SRR31800874-SRR31800877), Liao et al. 2022 (GSE195790), Descalzi et al. 2017 (GSE91396), Erdei et al. 2025 (PRJNA1137110) and the uninjured baseline (GSE73419).

## Code availability

Analysis and figure-generation code is available at https://github.com/SJRussell/Xeno_Brain_Regen (public on publication; available to editors and referees on request).

## Acknowledgments

The authors thank E. Switzer for animal husbandry and maintenance, and R. Colon and J. Mandal for laboratory assistance. We thank Jack Devine, David Shire, Eugene Mosharov and Martin Picard of the Mitochondrial Psychobiology Group for valuable discussions, Mijail Serruya for directing us to relevant literature and Nicolas Rouleau for editorial assistance. N.J.M. acknowledges support from the NSERC Discovery Grant (RGPIN-2021-03783). M.L. acknowledges support from the NIH (R01 AR005593 and R01 AR061988) and the W.M. Keck Foundation (5903). D.L.K. acknowledges support from the NIH (P41EB027062).

## Author contributions

N.J.M., D.L.K. and M.L. conceived the original experiment. N.J.M. and K.A.M. performed the animal experiments and collected tissue. S.J.R. and N.J.M. designed the analyses reported here. S.J.R. performed the bioinformatic analyses and prepared the figures. S.J.R. and N.J.M. wrote the manuscript, and D.L.K. and M.L. edited it. N.J.M., D.L.K. and M.L. acquired funding. All authors read and approved the final manuscript.

---

**Supplementary Figure 1.**
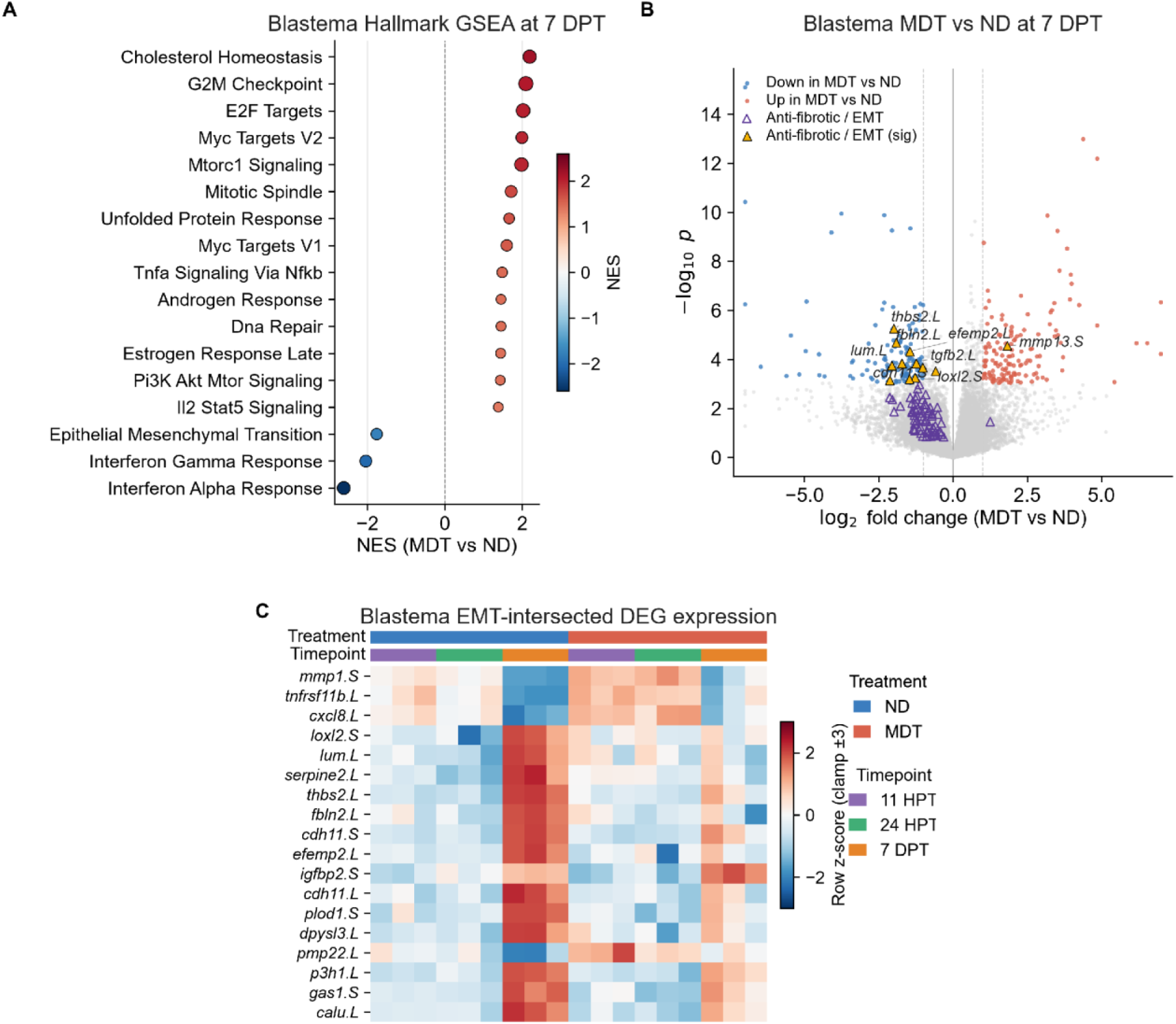
Blastema positive control: a pro-regenerative, anti-fibrotic state under treatment. **(A)** Blastema Hallmark GSEA at 7 DPT, MDT versus ND; NES on the x-axis and color, dot size reflects significance. **(B)** Blastema volcano plot, MDT versus ND at 7 DPT, with epithelial-mesenchymal transition (EMT) and anti-fibrotic genes overlaid as triangles (filled when significant). **(C)** Expression of EMT-intersected DEGs across conditions and timepoints (color, row z-score).

**Supplementary Figure 2.**
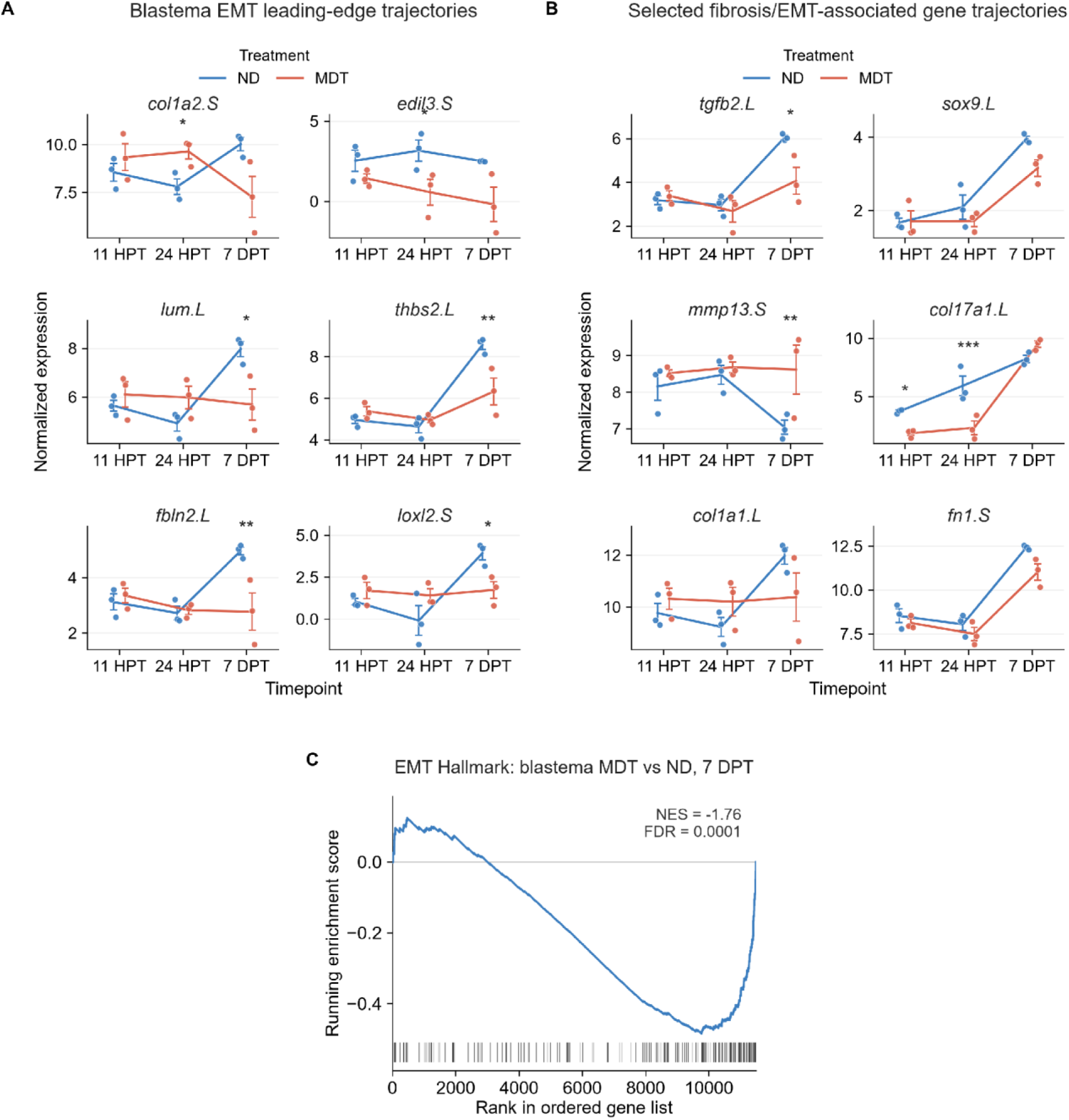
Extracellular matrix and fibrosis-associated expression in amputation-site tissue. **(A)** Leading-edge EMT gene trajectories; points are individual animals, lines connect group means and error bars show s.e.m. **(B)** Selected fibrosis and EMT-associated gene trajectories. **(C)** Running-enrichment (GSEA) plot for the EMT Hallmark, MDT versus ND at 7 DPT. In A and B, asterisks denote significant MDT-versus-ND differences.

**Supplementary Figure 3.**
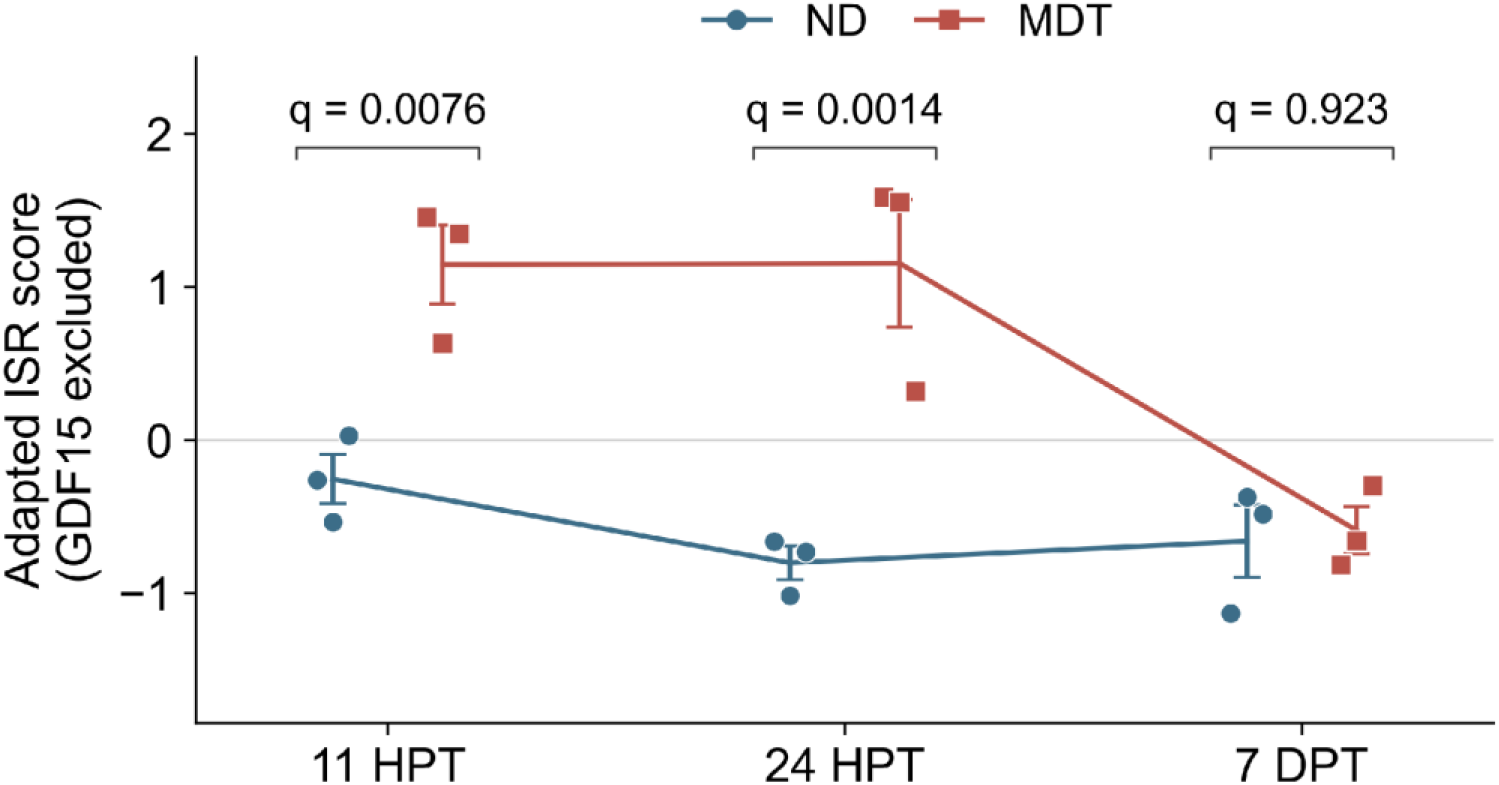
Early limb ISR-associated transcription under treatment. The exploratory *Xenopus*-adapted ISR score excludes GDF15 and uses 101 of 119 ISR genes. Points are individual animals (n=3 per treatment and timepoint); lines connect group means and error bars show s.e.m. MDT, multidrug treatment; ND, no device. Labels show BH-adjusted P values (q) for MDT versus ND from ordinary linear models of animal-level scores, corrected across the 119-test exploratory ISR family. The treatment difference decreased between 24 HPT and 7 DPT (interaction estimate=−1.88, 95% CI −2.95 to −0.82, q=0.010).

